# Beyond establishment: incorporating physiological performance into predictions of invasion risk

**DOI:** 10.64898/2026.08.28.747494

**Authors:** Lauren Vapillon, Soria Delva, María Bonafont Castelles, Jorge Assis, Diederik Strubbe, Tim Adriaens, Olivier De Clerck, Sofie Vranken

## Abstract

Biological invasions are a major driver of global change, reshaping ecosystems and threatening biodiversity worldwide. Anticipating where invaders will establish and where they will exert the strongest ecological impacts are key challenges for early detection and targeted management. Although Species Distribution Models (SDMs) are widely used to forecast biological invasions, they often provide uncertain estimates of establishment ranges and limited insight into invader performance, making it difficult to anticipate ecological impacts. Here, we address these limitations by integrating physiological information on invader performance with SDMs to identify regions of high invasion risk. Using the brown alga *Rugulopteryx okamurae,* one of the most prominent marine invaders in Europe, we first test alternative hypotheses of northern establishment limits: (i) a cold-survival constraint driven by winter temperatures and (ii) a growth constraint derived from the species’ thermal performance. To identify the more likely scenario, we combine cold-tolerance experiments with seasonal growth comparisons between the invader and a native macroalga *Dictyota dichotoma*, whose established distribution allows physiological performance to be directly related to realised presence. Finally, we project seasonal growth of the invader across the predicted establishment range as a proxy for biomass accumulation and potential ecological impacts. Our results indicate that northern limit in Europe will be more likely constrained by winter survival rather than growth, extending the potential establishment range of *Rugulopteryx* to mid-Norway. In contrast, the highest impacts are likely to remain concentrated in southern Europe, where thermal conditions sustain high year-round growth. Overall, our approach illustrates how understanding the physiological response of invaders to their environment can improve the interpretation of SDM outputs and help identify areas at greatest risk of impact within their potential establishment range.

## 1. Introduction

Invasive species pose a major threat to biodiversity and ecosystem services worldwide (Jaureguiberry et al., 2022; IPBES 2023; Haubrock et al., 2026) They displace native species, introduce diseases, and alter species interactions, all of which, in turn, can cascade to disrupt food webs and entire ecosystems (Blackburn et al., 2019; Ehrenfeld, 2010; Simberloff et al., 2013; Wainright et al., n.d.). Identifying and prioritising areas at high risk of invasion is therefore crucial to establish preventive measures and guide policy action (Guisan et al., 2013; Jiménez-Valverde et al., 2011). However, the geographic range in which species can establish is not necessarily equivalent to the range in which they will reach high abundance (O’Neill et al., 2021). This distinction is important as ecological impact generally increases with abundance rather than presence alone (Parker et al., 1999). Predicting invasion risk, therefore, requires a thorough understanding of the environmental conditions that govern not only the establishment of invasive species but also the processes that determine their abundance and the subsequent magnitude of their ecological impacts.

In ectotherms, environmental temperature is considered one of the main factors determining both geographic ranges and individual performance owing to its direct effect on cellular processes (Hochachka and Somero, 2002; Clarke, 2003; Pinsky et al., 2019; Sunday et al., 2012). Geographic distributions are often constrained by thermal thresholds acting on particular traits, such as survival (lethal limits), completion of the life cycle (reproductive limits), and growth requirements (e.g., Lüning, 1990; Andersen et al., 2015; Parratt et al., 2021). Such fitness-related traits often show large differences in thermal requirements (Kellermann et al., 2019; Delva et al., 2026), which underscores the need to identify which particular trait constrains the distribution of a given invasive species, and its associated thermal thresholds, to accurately predict the species’ potential establishment range.

Within the predicted establishment range, spatial and temporal variation in temperature further influences individual performance and biomass accumulation, as the thermal window for high performance is typically narrower than that for persistence (Bozinovic et al., 2020). This, in turn, creates temperature-driven variation in the ecological impact of invasive alien species. For example, several meta-analyses on aquatic invaders found higher ecological impacts in locations where temperatures closely match the species’ thermal growth optima or a specific part of their native thermal niche (Iacarella et al., 2015; Bennett et al., 2021). Despite growing evidence that thermal physiology shapes the invasion success of introduced ectotherms (Kelley, 2014), studies combining predictions of establishment ranges, and their thermal limits, with spatial predictions of thermal performance remain relatively rare (Briscoe Runquist et al., 2019; Fenollosa et al., 2025).

Establishment ranges are most commonly estimated using species distribution models (SDMs; e.g., Jeschke & Strayer, 2008; Jiménez-Valverde et al., 2011; Srivastava et al., 2019) which use the realized distribution of a species to infer its environmental requirements and predict suitable habitat (Elith & Leathwick, 2009). Yet, correlative SDMs do not explicitly capture the biological mechanisms constraining species’ distributions (Buckley et al., 2010) and they often perform poorly in predicting local abundance or performance across geographic ranges (Waldock et al., 2022; Lee-Yaw et al., 2022) To overcome these limitations, physiological information has increasingly been combined with correlative SDMs to identify the mechanistic basis of species’ range limits and to improve predictions of establishment and performance (Kearney & Porter, 2009; Elith et al., 2010; Claunch et al., 2023; Fenollosa et al., 2025). Thermal performance curves (TPCs) are particularly useful in this context, because they quantify how temperature influences fitness-related traits across the full thermal range experienced by a species (Huey & Stevenson, 1979; Angilletta, 2009). Because ecological impacts commonly increase with population abundance and biomass (Parker et al., 1999; Pearson et al., 2016; Bradley et al., 2019), thermal performance curves describing traits related to these processes can identify the temperatures that support high performance and, consequently, the regions where ecological impacts are most likely to arise.

Realising this potential, however, requires translating lab-derived physiological information into spatial predictions. Such integration is challenging, as broad-scale temperature datasets rarely capture the microclimatic conditions organisms experience in the field (Potter et al., 2013; Chevalier et al., 2024). This makes it difficult to accurately link lab-derived thermal responses with macroclimatic temperature layers or to compare them with thermal relationships inferred from SDMs calibrated on those layers (Briscoe Runquist et al., 2019). One potential way to evaluate the ecological relevance of lab-derived thresholds is through comparisons with functionally equivalent closely related native species whose distributions and thermal ecology are well characterised. These species could then provide an ecological reference against which lab-derived thermal thresholds can be compared.

In this study, we focus on the invasive brown seaweed *Rugulopteryx okamurae* (E.Y.Dawson) I.K.Hwang, W.J.Lee & H.S.Kim 2009 (hereafter *Rugulopteryx*), one of the most prominent invaders in Europe, and compare its thermal ecology with that of the closely related native species *Dictyota dichotoma* (Hudson) J.V.Lamouroux 1809 (hereafter *Dictyota). Dictyota* is morphologically very similar to *Rugulopteryx* and has been extensively studied, with a particular focus on its thermal performance (Bogaert et al., 2016; Delva et al., 2023; Delva et al., 2026). Importantly, the species is one of the most widely distributed species of its genus in Europe (Tronholm et al., 2010), making it a useful reference for evaluating the establishment and performance potential of *Rugulopteryx* across Europe.

Originally introduced in Europe in the Thau Lagoon (France; Verlaque et al., 2009), *Rugulopteryx* began to display invasive behaviour following its introduction in the Strait of Gibraltar in 2015 (Ocaña et al., 2016). From there, the species rapidly expanded its range across the western Mediterranean Sea and adjacent Atlantic regions, with recent records demonstrating a continued northward expansion (Díaz-Tapia et al., 2025). *Rugulopteryx* displays strong competition and colonisation abilities, forming dense mats that completely transform benthic assemblages by displacing species, homogenising communities and altering the physicochemical environment (Borriglione et al., 2024). Owing to these impacts, *Rugulopteryx* has been included on the List of Invasive Alien Species of Union concern, enforcing member states to set up surveillance, implement preventative pathway action plans and adopt measures to mitigate adverse impact and restore invaded areas (European Union (EU), 2022). Despite evidence of negative impacts of several other non-native seaweeds, it remains the only macroalga listed so far (van der Loos et al., 2024).

Here, we integrate species distribution modelling with physiological experiments to assess the thermal mechanisms governing the establishment and performance of *Rugulopteryx* across Europe, using the native *Dictyota* as an ecological reference. First, we quantified thermal responses of growth and survival for both *Rugulopteryx* and *Dictyota* to characterise species-specific responses to temperature. To predict the establishment range for *Rugulopteryx* in Europe, we constructed two alternative SDMs representing competing hypotheses of establishment range restriction: a traditional correlative SDM in which the species’ range is constrained by cold winter temperatures (i.e., a lethal limit; *sensu* Breeman, 1988) and a hybrid SDM in which the species is constrained by a temperature-dependent growth limit. Then, we projected seasonal growth performance across the predicted establishment range to assess spatial variation in performance and invasion potential across Europe. By identifying regions where *Rugulopteryx* is most likely to establish as well as the part of this area where it is most likely to effectively exert ecological impacts, we provide a mechanistic basis for focusing surveillance and management efforts.

## 2. Material & Methods

### 2.1 Thermal performance experiment

To obtain thermal performance curves for growth, *Rugulopteryx* individuals were collected from Marseille, France (43°17’5.8”N, 5°20’58.7”E) in January 2021. Cultures were directly established from the sampled adult thalli, as no fertile individuals were observed. To this end, we treated algal individuals with GeO_2_ and the broad-spectrum antibiotic streptomycin to reduce diatom and bacterial contamination, respectively. To further limit contamination, cultures were initially maintained under low nutrient conditions through regular addition of modified Provasoli medium (West and McBride, 1999). When no signs of contamination were observed, individuals were further grown in crystallizing dishes filled with autoclaved natural seawater enriched with 10 mL L^-1^ modified Provasoli medium (hereafter enriched seawater). For *Dictyota*, we used laboratory-reared gametophytes derived from fertile sporophytes collected in August 2018 in Île de Noirmoutier, France (47°01’37.2”N, 2°18’18.0”W). These individuals were obtained according to the procedures outlined in Delva et al. (2026), which are based on the isolation and subsequent development of spores. All individuals of *Dictyota* and *Rugulopteryx* were maintained in enriched seawater under a 12h:12h light:dark cycle for several months prior to the experiments. Individuals were selected to represent the most ecologically relevant life stages for the study region. Specifically, gametophytes were used for *Dictyota*, reflecting their higher frequency in populations along the northeastern Atlantic (Vranken et al., 2026). For *Rugulopteryx*, the life cycle phase of the invasive individuals could not be confirmed due to the absence of reproductive structures and the isomorphic nature of the species. Nevertheless, the material represents the adult vegetative thallus, which is the functionally dominant form in invasive populations. The presence of (mono)spores and the lack of observed gametes in monitored *Rugulopteryx* populations suggest that the sexual cycle is rarely completed and that these invasive populations may be largely maintained by sporophytes (Rosas-Guerrero et al., 2026; Santos et al., 2026).

To determine the thermal growth curve of both *Rugulopteryx* and *Dictyota*, clonal apices from four individuals per species were exposed to a gradient of seven temperatures (11°C, 16°C, 20°C, 22°C, 24°C, 27°C, and 30°C), with one apex per individual assigned to each temperature (n = 28 apices per species). These assay temperatures were selected to cover nearly the entire range of temperatures experienced across the current invaded range of *Rugulopteryx* (Román & Vázquez, 2025), while also allowing to construct a reliable thermal growth curve for *Dictyota* (see Delva et al., 2023, 2026). The experiment was conducted in thermal incubators and temperature was monitored continuously using thermal loggers (Thermochron iButton DS1921G-F5#, Maxim Integrated, United States). Each algal apex was grown in a separate crystallizing dish filled with 30 mL of enriched seawater medium under a light intensity of 16-27 µmol m^-2^ s^-1^ (cool white LEDs, Ohmeron, Belgium) and a 12h:12h light:dark cycle. At the start of the experiment, all apices were gradually acclimated to their respective temperatures using a thermal ramping of 2°C per half hour. The experiment lasted for 14 days, and the medium was refreshed after 7 days. We photographed each algal apex at the beginning and at the end of the experiment and quantified their surface area using the image analysis software ImageJ (Schindelin et al., 2015). Relative growth rate (RGR) was then calculated according to the following equation:

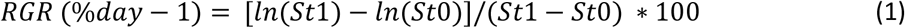

where St_0_ and St_1_ represent the surface area at the beginning (t_0_) and at the end (t_1_) of the experiment. Negative growth rates were converted to zero following Delva et al. (2023, 2026).

Following (Angilletta, 2006) we used an information theoretic approach to select the thermal model that described the data well without overfitting. Based on these analyses, the model described by Lobry et al. (1991) was chosen to fit the growth performance curve of both *Rugulopteryx* and *Dictyota*:

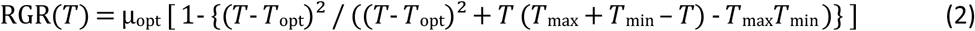

where *T* is the temperature (°C), µ_opt_ is the maximum growth rate (% day^-1^) at the optimum temperature *T*_opt_ (°C), and *T_max_* and *T_min_* are the minimum and maximum temperatures for growth respectively (°C). The model was fitted using non-linear least squares regression, employing the *nlsLM* function of the R package *minpack.lm* (Elzhov et al., 2026). To assess whether *Dictyota and Rugulopteryx* differed in their thermal performance, several performance metrics were derived from the fitted curve, e. i. the thermal performance breadth (measured as the thermal range across which the growth rate was at least 80% of the maximum growth; see Eggert, 2012), and the lower and upper thermal limits of the performance breadth.

### 2.2 Cold tolerance experiment

A lab experiment was performed to directly assess the cold survival limit of *Rugulopteryx* and inform our SDM modelling on the northern range limit. To provide a comparative ecological baseline, we assessed the cold tolerance of *Dictyota* using the same experimental approach. *Rugulopteryx* individuals were collected from Marbella, Spain (36°30’5.4”N, 4°50’23.5”W), while those from Dictyota were collected from Goes, Netherlands (51°35’22.3”N, 3°31’53.2”E). Prior to the experiment, individuals were grown under laboratory conditions in 1-L bottles filled with enriched seawater and aeration. Cultures were maintained under a 12h:12h light:dark cycle at 17°C and a light intensity of 15 µmol m^-2^ s^-1^, and the growth medium was refreshed weekly. Both species were kept under these conditions for at least 24 months before the experiment, erasing putative imprints of the natural environment.

To assess cold survival limits, five clonal apices of each individual were exposed to a stepwise cooling starting from 12°C. Temperature was decreased by 2°C every two weeks until reaching 4°C, corresponding to the lowest temperature experienced by *Rugulopteryx* in its native range (Fig 4). After reaching 4°C, individuals were maintained at this temperature for a prolonged period to simulate winter conditions along the northern European coastlines. Throughout the experiment, each algal apex was maintained in a separate crystallizing dish containing 60 mL of enriched seawater medium under a 12h:12h light:dark cycle and a light intensity of 5 µmol m^-2^ s^-1^. Algal health and survival were monitored biweekly by measuring the maximum quantum yield of photosystem II (F_v_/F_m_) using a MAXI-imaging-PAM M-series (Heinz Walz GmbH) after 15 minutes of dark adaptation, and by visual assessment of tissue necrosis. The culture medium was refreshed after each measurement.

### 2.3 Occurrence records and environmental predictors

Georeferenced *Rugulopteryx* occurrences were compiled from van der Loos et al., 2024 (available from https://zenodo.org/records/7798640) supplemented with more recently published distribution records and manually curated distribution records from iNaturalist (available from https://www.inaturalist.org) and GBIF (available from https://www.gbif.org/what-is-gbif). All records were carefully reviewed, and those with doubtful identifications or uncertain geographic locations were excluded (e.g., GBIF records from Australia and New Zealand, as well as records from northern Taiwan and the south-east coast of China). This resulted in a dataset of 823 verified occurrences spanning the Northwest Pacific (i.e., the native range), the Mediterranean Sea, Macaronesia and the Northeast Atlantic (i.e., the invasive range of the species). Data were further filtered to present-day conditions (2000–2020), and inland points less than 25 km from the coast were relocated to the nearest coastal cell. To reflect the vertical distribution of the species, occurrences were restricted to a maximum depth of 50 m (García-Gómez et al., 2020, Estévez et al. 2022). Following this procedure, a final dataset of 248 native and 221 non-native occurrence records was assembled.

A set of environmental predictors was selected based on the biological requirements of *Rugulopteryx,* and variables commonly used to model the distribution of intertidal and subtidal brown macroalgae (Sainz-Villegas et al., 2022, Delva et al. 2026). Long-term maximum (LtMax) and minimum (LtMin) ocean temperatures, LtMin salinity, and mean nitrate concentrations were obtained from Bio-ORACLE v3.0 benthic layers (Assis et al., 2024)for present-day conditions (2000–2020) at a 0.05° resolution. Mean photosynthetically active radiation (PAR) was acquired from bio-ORACLE v2.0 (Assis, Tyberghein, et al., 2018) and aligned to the v3.0 resolution. In addition, an annual growth rate (AGR) predictor was developed from the species’ thermal growth response to include in the hybrid SDM. To this end, monthly ocean temperatures were downloaded from the Copernicus Marine Service (CMEMS; Product ID: GLOBAL_MULTIYEAR_PHY_001_030; https://data.marine.copernicus.eu/products) at a temporal and spatial resolution consistent with the Bio-ORACLE v3.0 layers. First, monthly growth rates were estimated by projecting the thermal growth curve described in Section 2.1 onto the monthly CMEMS layers, with negative growth values set to zero. These were then averaged to generate a single AGR predictor representing mean growth over the year. To constrain predictions within the species’ bathymetric limits, all predictors were restricted to the maximum occurrence depth of the species (50 m), augmented with a 15m depth buffer. The spatial extent of predictors was further limited to: (i) the species’ native range, defined by the area of native occurrences extended by 12° in all directions, and (ii) the potential invaded range defined as the European coastlines, and bounded by coordinates Lat (−30°, 40°) and Lon (25°, 70°). Prior to modelling, Pearson’s correlation coefficient was estimated between each pair of predictors.

### 2.4 Pseudo-absence selection

To account for the lack of absence records in the compiled dataset, pseudo-absences (p-a) were generated in a 1:1 ratio with occurrence data (see below, (Barbet-Massin et al., 2012). Following (Chapman et al., 2019), we defined the background area for p-a sampling to account for the potential non-equilibrium dynamics of invasive species undergoing rapid range expansion. This approach prevents sampling p-a in unoccupied regions that may be environmentally suitable but have not yet been reached by the invader due to dispersal limitations. The background domain combined: (i) the accessible area, defined as regions where the invader has had opportunity to disperse but has not established, and (ii) the unsuitable area, reflecting known biological constraints on the species’ establishment. In the non-native range, the accessible area was delineated as ecoregions where at least one *Rugulopteryx* occurrence was recorded. Unsuitable areas were defined as regions with depths greater than 50 m (i.e., the maximum depth of the species) or with ocean temperatures exceeding 28.5 °C (i.e., the LtMax temperature experienced by *Rugulopteryx* in its native range). Unsuitable areas were restricted to neighbouring ecoregions only. In the native range, where the species is considered at equilibrium with its environment, the background domain was defined exclusively as the accessible area, delineated as all ecoregions with at least one recorded *Rugulopteryx* occurrence, together with all neighbouring ecoregions. An initial set of p-a of ten times the occurrence count was generated in the background domain, ensuring a minimum distance of 100km between p-a and any occurrence point, and 5.6 km (0.05° cell resolution) between every p-a. p–a sampling was weighted using a seaweed bias grid assembled from the Global Biodiversity Information Facility (GBIF) and representing the log-transformed number of records of Phaeophyceae, Rhodophyceae, and Ulvophyceae per grid cell. Higher grid values increase p-a sampling probability in cells with greater macroalgal sampling effort, where the absence of *Rugulopteryx* can therefore be inferred with higher confidence. Final p-a were extracted at a 1:1 ratio with occurrence data by generating environmentally dissimilar clusters. Specifically, K-means clustering was applied to the environmental data of candidate p-a points, using the number of occurrence records as the *K* parameter. This procedure accounted for spatial sampling bias and minimized redundancy in environmental information.

### 2.5 Species Distribution Modelling (SDM)

To project the potential establishment range of *Rugulopteryx* in Europe, and particularly to identify its most likely northern distribution limit, we developed two SDMs representing alternative hypotheses of range limitation. First, a correlative SDM assumed that establishment at the northern limit is primarily constrained by cold winter temperatures (i.e., survival limit). This model included the five environmental variables listed above (i.e. LtMin and LtMax temperature, LtMin salinity, mean nitrate concentration and mean PAR. Among these, LtMin temperature (i.e., the temperature of the coldest month, on average) primarily reflects survival constraints at the northern limit of the predicted distribution. Although temperature and AGR predictors were strongly correlated (Pearson’s r = 0.72 between LtMax and LtMin temperature, and r = 0.80 between LtMax temperature and AGR; Fig. S1), all variables were retained for modelling, as all three predictors represent distinct ecological constraints acting at different margins of the species’ distribution (Dormann et al., 2013). Second, a hybrid SDM assumed that northern establishment is limited by growth performance and replaced LtMin temperature with the AGR predictor derived from the species’ thermal growth response (see above). All other predictors were identical between models. Both SDMs were constructed using the boosted regression trees (BRT) machine learning algorithm as it previously achieved the best accuracy for modelling *Rugulopteryx* distribution among three other statistical models (Sainz-Villegas et al., 2022) and efficiently handles high collinearity between predictors (Elith et al., 2008). Models were trained on pooled occurrence data from both the native and the non-native ranges to better capture the species’ full environmental niche, a common approach in invasion biology that enhances predictive performances in invaded regions (Broennimann & Guisan, 2008, Elith 2017). To ensure a single record per gridded cell, occurrence data were resampled to the 0.05° spatial resolution of bio-ORACLE v3.0 layers, resulting in a final dataset of 207 presences (122 native and 85 non-native) and an equal number of pseudo-absences. To minimize overoptimistic assessment of model predictive performances that may arise from spatial autocorrelation (Ploton et al., 2020), we conducted cross-validation by partitioning the data into spatial blocks and randomly distributing these blocks across 10 folds. Block size was estimated using the *cv_spatial_autocor* function of the blockCV R Package (Valavi et al., 2019), or by dividing the longitudinal extent of the data by the number of folds, depending on which method yielded the smallest block size. Optimal BRT model parameters were selected by training models with combinations of trees (50–2000 in steps of 50), tree complexity (1–4), and learning rate (0.01, 0.005 and 0.001) across nine folds of data, while one independent fold was withheld at a time to assess predictive performance. Monotonicity constraints were applied to all predictors (positive for all except long-term maximum ocean temperature) to reduce overfitting and enhance transferability of the models (Hofner et al., 2011). The models were evaluated based on the True skill statistic (TSS) (TSS > 0.8 indicating high accuracy, Allouche et al., 2006) and the area under the receiver operating characteristic curve (AUC) (values near 1 signifying alignment with the observed data, Hirzel et al., 2006). Final models were run using the combination of parameters that achieved the highest mean TSS across all cross-validation folds. Finally, each model type was run five times, with a new set of pseudo-absences sampled at each iteration. Suitability estimates are reported as the median across model replicates and presented as continuous likelihood probabilities ranging from 0 to 1. Corresponding binary presence–absence maps were generated using a threshold that maximized sensitivity and specificity (Assis et al., 2018; Assis et al., 2016). These reclassified outputs should be interpreted cautiously, as such threshold-based approaches may result in more conservative estimates of suitable area, particularly in the non-native range.

### 2.6 Spatial projections of seasonal growth

To visualise spatial variation in seasonal growth and estimate putative ecological impact, the thermal growth response of *Rugulopteryx* was projected across 10 representative locations spanning the full thermal gradient of its native, invasive and projected establishment range. In the native range this was Shibushi, Gangneung, and Sapporo. In the non-native range, three sites were selected from the projected establishment range (Quiberon, The Hague and Bergen) and four from the current distribution. The latter were chosen to contrast areas experiencing severe invasions (i.e. Ceuta, Marseille) with recently colonised regions where establishment remains at an early stage (i.e. Ariceales, Bilbao) (Fig.6). For each location, *Rugulopteryx* thermal growth curve was projected onto monthly ocean temperatures (CMEMS; see above), and negative values were set to zero, yielding monthly estimates of growth. These projections capture the seasonal thermal favourability for growth, integrating both the duration (seasonality) and magnitude (intensity) of growth. Spatial projections of seasonal growth were used for two complementary purposes. First, projected seasonal growth at sites within the potential establishment range was used, in combination with the cold-tolerance experiment, to evaluate physiological constraints on northern establishment and support the interpretation of SDM outputs. Comparable projections were generated for the native *Dictyota,* whose realised distribution provides an ecological benchmark for linking physiological performance to establishment outcomes. Second, seasonal growth projections were used to identify areas most likely to experience high ecological impact within the potential establishment range of *Rugulopteryx*.

## 3. Results

### 3.1 Thermal growth response

Thermal growth responses differed considerably between *Rugulopteryx* and *Dictyota*, especially at higher experimental temperatures (Fig. 3A) as shown by their maximum growth rate and optimum temperature. Maximum growth rates were estimated at 16.4 % day⁻¹ (CI: 15.3–17.4 % day⁻¹) for *Rugulopteryx*, at an optimal temperature of 26.5 °C whereas *Dictyota* reached a maximum growth rate of 12.07 % day⁻¹ (CI: 10.0–14.2 % day⁻¹) at an optimal temperature of 22.4 °C (Fig. 3A). While *Rugulopteryx* displayed a slightly narrower thermal performance breadth than *Dictyota* (8.0°C and 9.1°C respectively), this difference was not significant. However, both lower and upper performance limits were significantly higher for *Rugulopteryx* (L: 20.9 °C, U:to 28.9 °C) compared to *Dictyota* (L: 17.2 °C, U: 26.4 °C). At the highest experimental temperature of 30°C, all *Dictyota* individuals showed negative growth rates (here converted to zero), while the majority of *Rugulopteryx* individuals still displayed positive growth. Conversely, at suboptimal temperatures, growth rates of *Rugulopteryx* were similar to those of *Dictyota*. For example, at 11 °C, both species were growing at reduced rates (*Rugulopteryx*: 3.7 % day⁻¹, *Dictyota:* 2.6 % day⁻¹), corresponding to approximately 22 % and 21 % of their respective maximum growth rates.

**Fig.1.**
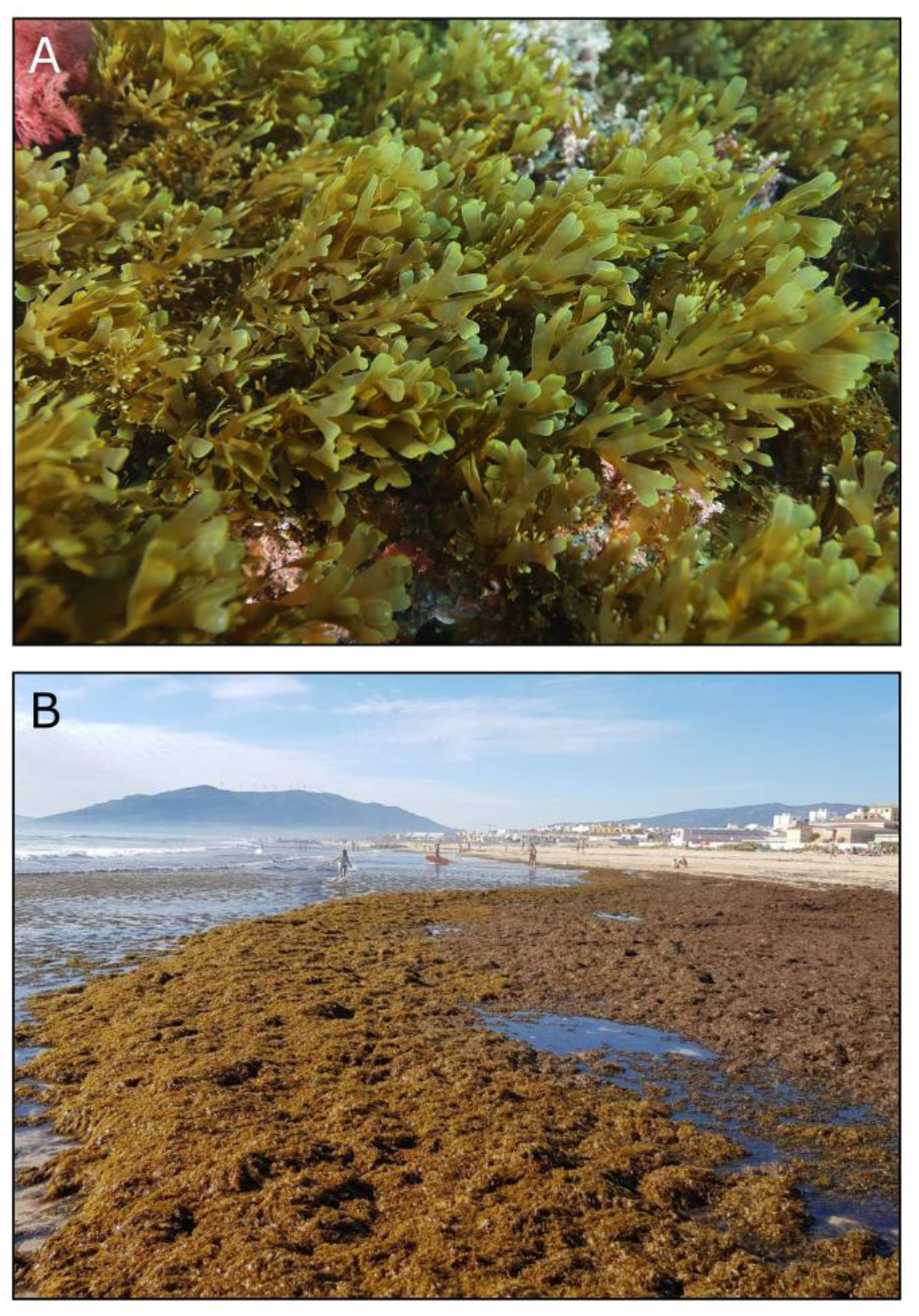
Photographs of A) underwater stand of Rugulopteryx in the Marseille area (©Sandrine Ruitton) and B) Rugulopteryx beach cast (Tarifa, Spain 2025, ©María Altamirano).

**Fig.2.**
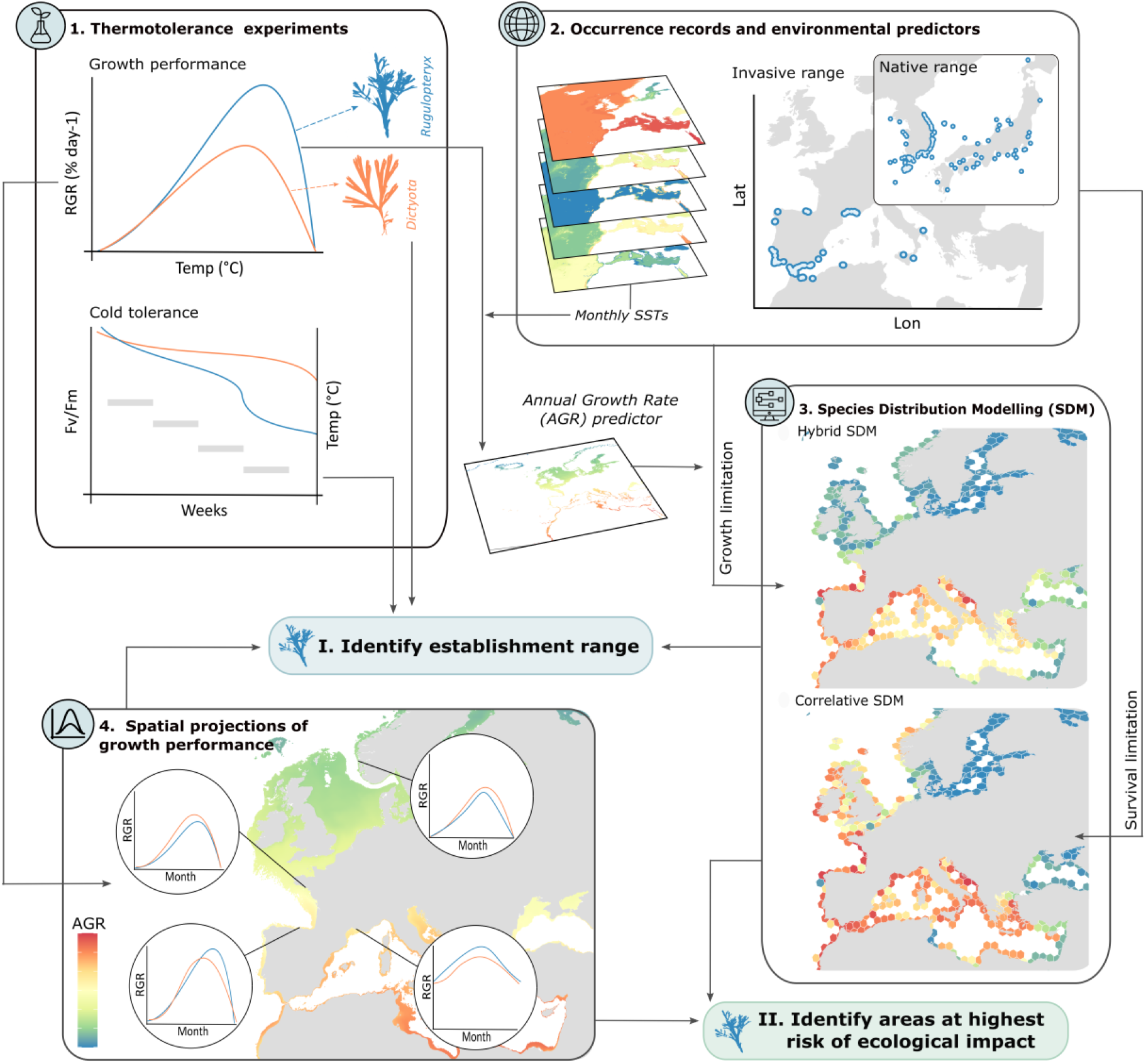
Illustration of the study workflow.

**Fig. 3.**
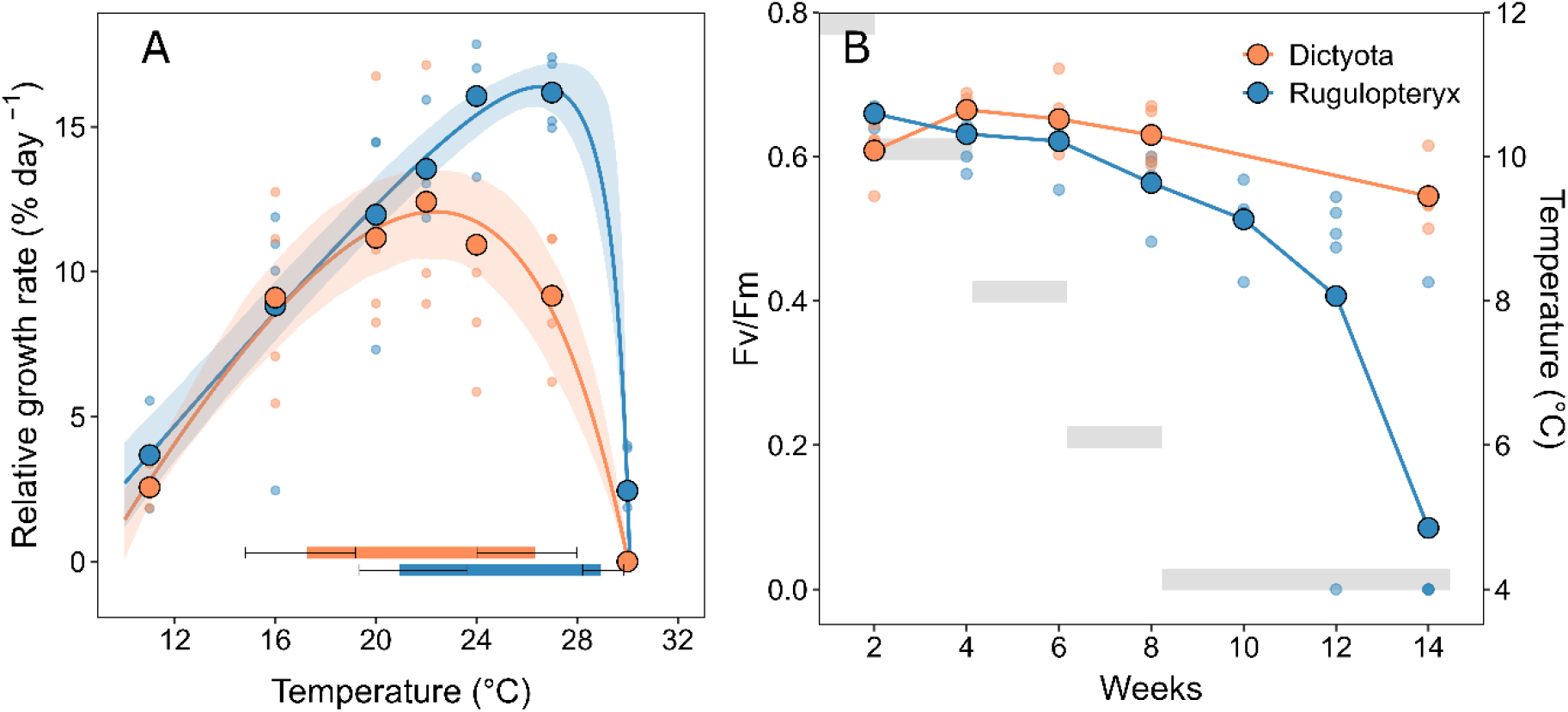
A) Thermal performance curve for relative growth rate (%day^-1^) for Rugulopteryx (blue) and Dictyota dichotoma (orange). The small dots represent the observed values, and the larger ones the mean across replicates. The lines depict the thermal response curve described by the Lobry model (Lobry et al. 1991), while the shaded areas represent the 95% confidence intervals obtained by bootstrap resampling (n bootstrap = 1000). Horizontal bars represent the thermal performance breath of each species while error bars indicate the 95% confidence intervals obtained by bootstrap resampling (n bootstrap = 1000). B) Fv/Fm values for Rugulopteryx (blue) and Dictyota dichotoma (orange) individuals exposed to 12°C, 10°C, 8°C, and 6°C for 2 weeks each, and to 4°C for 6 consecutive weeks. The grey blocks represent the temperature change over time.

### 3.2 Cold tolerance experiment

*Rugulopteryx* displayed sustained photosynthetic activity (*F*_v_/*F*_m_) across decreasing temperatures, from 12 °C to 6 °C (Fig. 3B). However, *F*_v_/*F*_m_ values drastically declined at 4°C, with the first individual reaching an *F*_v_/*F*_m_ of zero after four weeks, indicating the complete cessation of photosynthetic activity. After six weeks at 4 °C, all individuals showed an *F*_v_/*F*_m_ of zero, although no necrosis was observed. In contrast, *Dictyota* maintained relatively high *F*_v_/*F*_m_ values throughout the experiment, indicating broad tolerance to the low temperatures tested (Fig. 3B).

### 3.3 Potential establishment range of *Rugulopteryx* in Europe

Both SDMs performed well, although the correlative model achieved higher predictive accuracy than the hybrid one (TSS = 0.87 ± 0.04, AUC = 0.94 ± 0.02 versus TSS = 0.76 ± 0.03, AUC = 0.88 ± 0.01), largely due to a higher specificity (Fig. S2). Among environmental predictors, PAR was the most influential, followed by temperature-related variables (long-term maximum and long-term minimum temperature or the derived annual growth rate (AGR)), whereas nitrate and salinity contributed comparatively little (Fig. S2).

The projected northern establishment limit of *Rugulopteryx* in Europe differed markedly between the two models (Fig. 5). Whereas the correlative SDM classified most northern European coastlines up to mid-Norway as highly to moderately suitable (Fig. 5A,C), the hybrid SDM identified the central Bay of Biscay as the northern boundary to establishment (Fig. 5B,D). Although model outputs are primarily interpreted as continuous likelihood, reclassifying suitability into presence–absence highlighted the magnitude of the divergence between models, revealing a shift of about 2,300 km in the projected northern establishment limit (Fig. 5C,D). This divergence reflects the different mechanisms constraining establishment in each model. In the native range, the lowest long-term minimum temperature experienced by the species is 4.2°C (Fig. 4), a threshold closely captured by the correlative SDM as likelihood suitability declines below 0.5 where winter temperatures fall beneath this value. This pattern is observed not only along the Norwegian coastline but also along the Danish and German coasts of the North Sea, where similar thermal regimes occur (Fig. 4, Fig. 5A,C). In contrast, under the hybrid SDM, suitability starts to decline sharply north of the Bay of Biscay and AGR falls below ∼5.5% day⁻¹ (i.e., the lowest value observed within the native range) north of southern England (Fig. 4, Fig. 5B,D). Importantly, occurrences recorded near the northern edge of the native distribution simultaneously experience the lowest winter temperatures and the lowest AGRs projected (Fig. 4). While these two constraints closely covary in the native range, they become decoupled in the invasive range, where large areas of northern Europe that exhibit low AGRs (AGR < 5.5% day⁻¹) are areas where winter temperatures remain far above 4°C (Fig. 4).

**Fig. 4.**
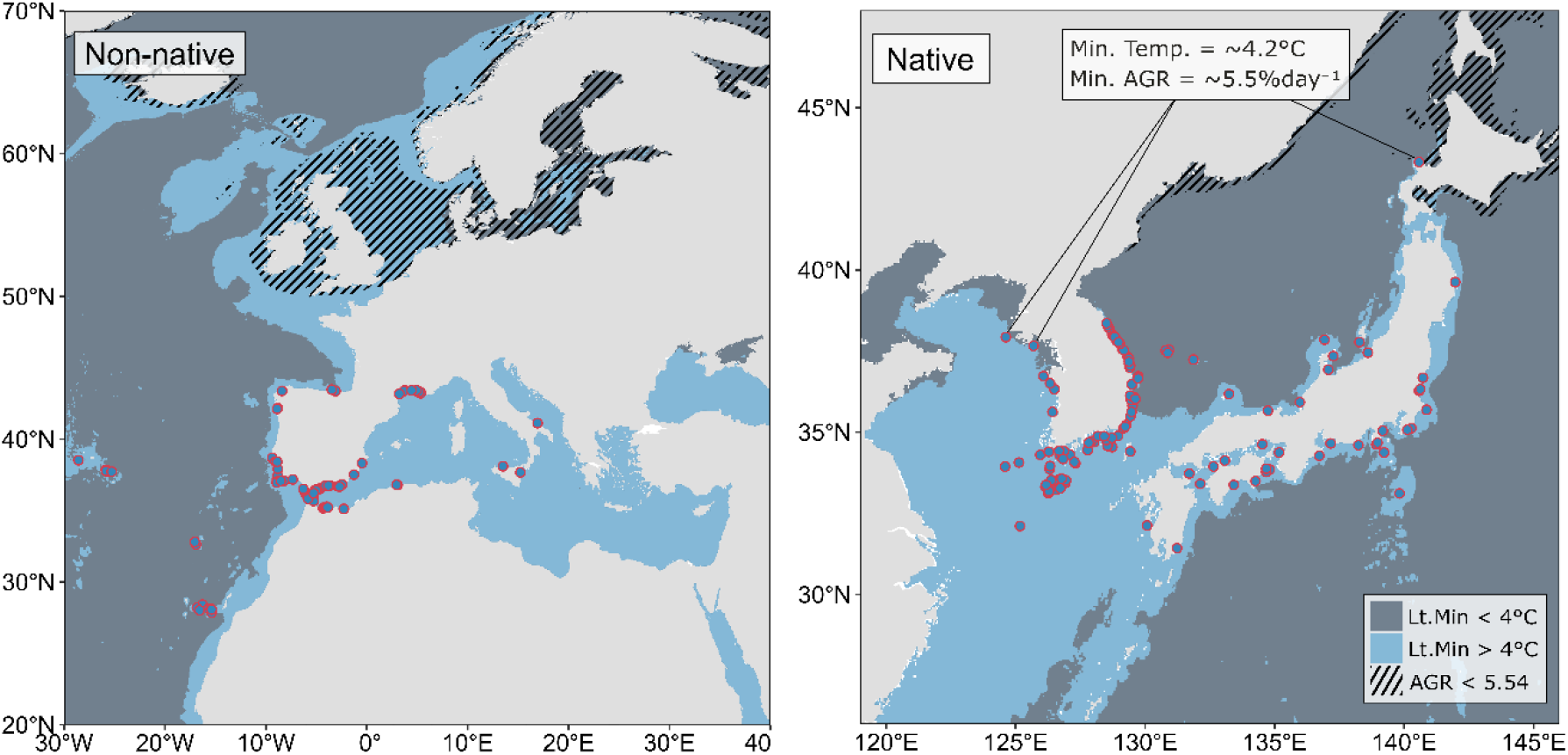
Native and Non-native Rugulopteryx occurrence records used to train the machine learning algorithms. Long-term minimum ocean temperature mask in the non-native and native range of the species. Dashed area represent annual growth rates (AGR) below 5.5% day-1 (i.e., the lowest value observed within the native range).

**Fig. 5.**
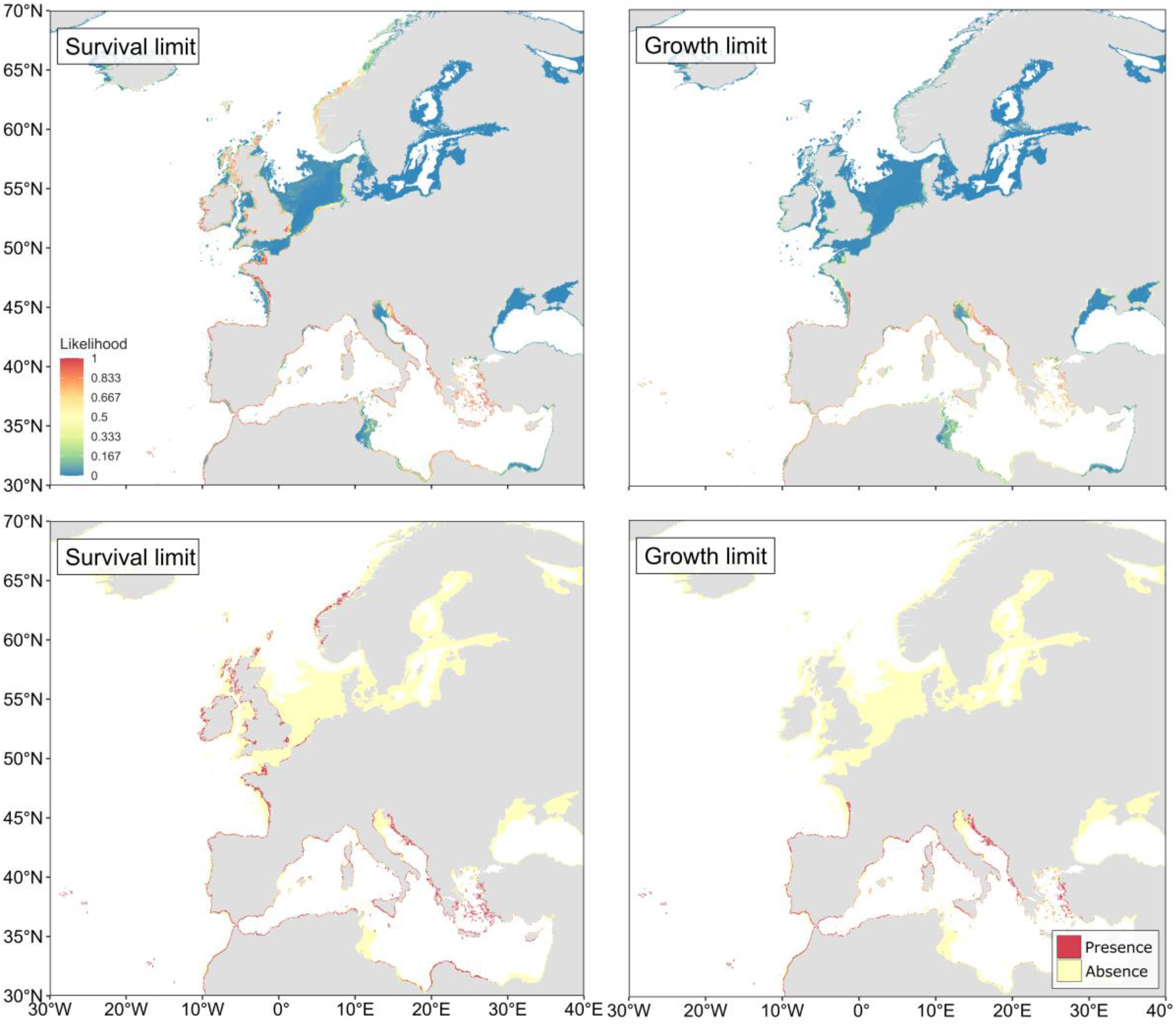
Habitat suitability for Rugulopteryx along European coastlines under present-day conditions, projected from the A) correlative and B) hybrid SDMs, along with their reclassified distributions C) and D). Reclassification thresholds are 0.73+-0.06 and 0.66+-0.08, respectively.

South of the Bay of Biscay, both SDMs predicted broadly similar suitability patterns (Fig. 5). Moderate to high suitability was consistently projected across most of the Mediterranean basin, including the Strait of Gibraltar, the Spanish coastline, the Balearic Islands, Sardinia, Sicily, and much of the Adriatic, Ionian, and Aegean Seas. Suitable conditions were also projected throughout Macaronesia (Azores, Madeira, and the Canary Islands), along the Moroccan coast, and up the Atlantic coast of the Iberian Peninsula to northern Spain, consistent with recent records from this region (Díaz-Tapia et al., 2025). In contrast, both models projected low suitability in the Gulf of Gabes, the eastern Levantine Sea, and the Red Sea. Additional unsuitable areas were consistently identified in the Black and Baltic Seas.

### 3.4 Spatial projections of seasonal growth in Europe

The projected growth of *Rugulopteryx* across Europe shows strong latitudinal variation in both the intensity and seasonality of growth (Fig. 6). They shift from high performance and low seasonality at the warmest edge of the invasive distribution, to reduced performance and marked seasonality at the coldest edge of the projected establishment range. North of the Bay of Biscay, in areas where the potential establishment ranges diverge, projected growth (see above) indicates active growth during spring and summer (approximately May to November), up to Bergen (Fig. 6). This pronounced seasonality is comparable to that observed at the centre and northern edge of the native distribution (i.e., Gangneung and Sapporo; Fig. S4), although peak growth there approaches maximum rates during the warmest months (> 85%; ∼14% day⁻¹) whereas it is predicted to remain more constrained in the invasive range, reaching ∼70% to ∼50% of maximum growth from Quiberon to Bergen (Fig. 6). These differences reflect a narrower range of annual sea temperature in northern Europe, where winter minima remain similar to the native range but summer temperatures are reduced (e.g. LtMax temperature in Sapporo: 24°C vs Bergen: 17°C; The Hague: 20°C; Fig. S6, Table S1). Comparing the seasonal growth of *Rugulopteryx* with that of the native *Dictyota* reveals highly similar patterns. From Quiberon up to Bergen, both species exhibit comparable seasonality (May to November) and growth intensity. This is particularly striking when comparing absolute rates (% day⁻¹), with both species projected to reach the same maximum absolute growth rates in August (e.g., Quiberon and The Hague: ∼13% day⁻¹, Bergen: ∼10% day⁻¹; Fig S5), which corresponds to approximately half of the maximum growth capacity of *Rugulopteryx* (50-60%), but nearly 80% to 95% of that of *Dictyota*, consistent with the lower thermal optimum and higher cold tolerance of the native species. This close similarity in seasonal growth suggests that thermal conditions in northern Europe may still support growth regimes compatible with the persistence of *Rugulopteryx* populations, as evidenced by the established distribution of *Dictyota*.

**Fig. 6.**
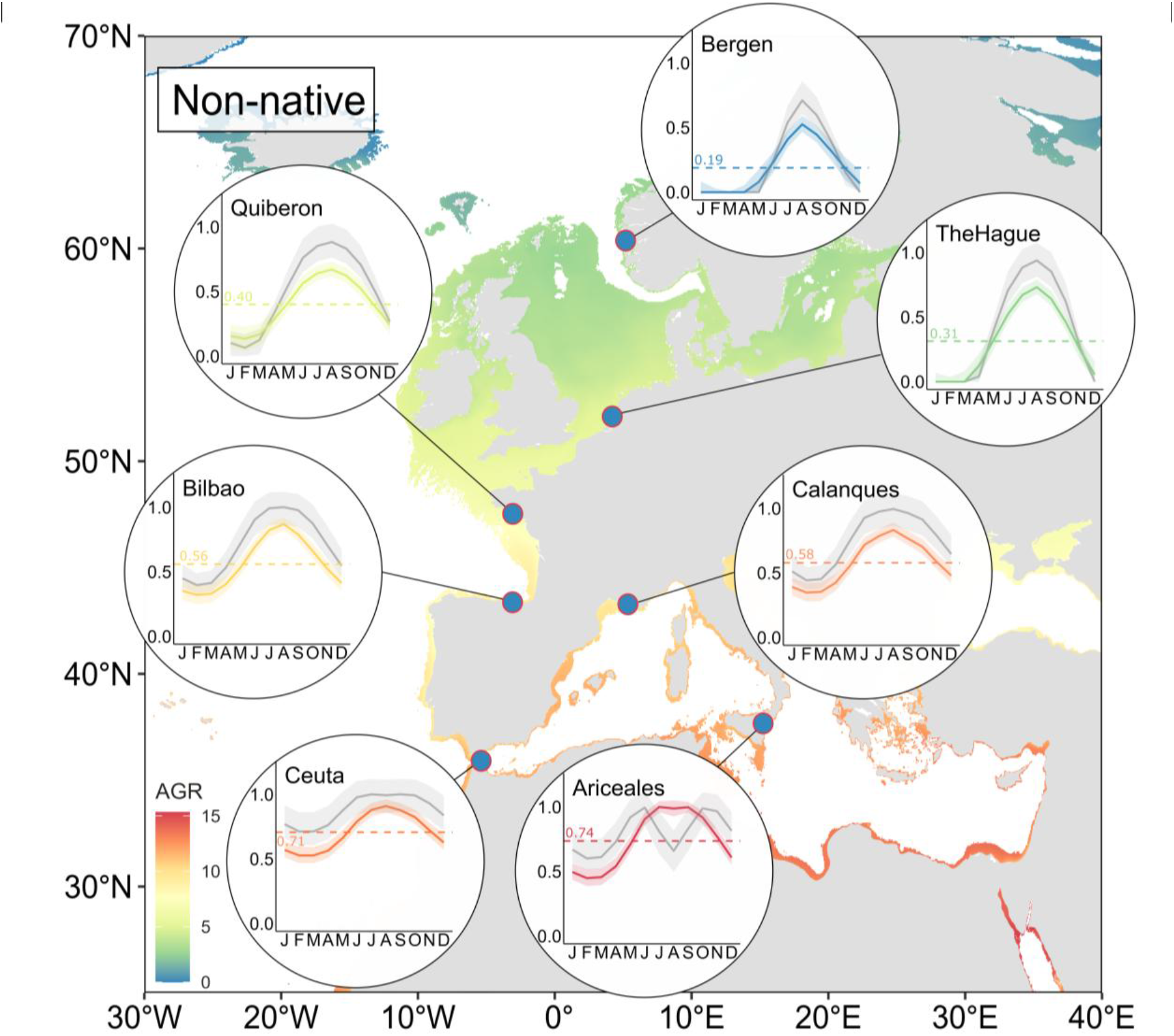
Annual growth rates projected in the Invasive range of Rugulopteryx. Individual line graphs depict the relative monthly growth rates of Rugulopteryx (coloured lines) and Dictyota (grey lines), standardized to the maximum predicted growth rate of each species (range: 0–1), as estimated by the fitted Lobry model (Lobry, 1991). Shaded ribbons represent the 95% confidence interval obtained by bootstrap resampling (n = 1000 bootstraps). Horizontal dashed lines depict the average annual growth rates of Rugulopteryx at each location.

At the warm edge of the invasive distribution (e.g. Ceuta and Ariceales), thermal conditions remain favourable throughout the year, with the species maintaining more than 50% of its maximum growth capacity in most months. These locations are characterised by high AGRs (e.g, Ceuta: 11.6% day⁻¹, Ariceales: 12.1% day⁻¹) corresponding to ∼70% of maximum growth. Such sustained high growth is also observed at the warm edge of the native range in Shibushi (Fig. S4). Across the Mediterranean basin, similar or higher AGRs (> 11 % day⁻¹) are consistently projected in regions classified as highly suitable by both SDMs (Fig. 5, Fig. 6). This pattern reflects a combination of high growth intensity and low seasonality, resulting in prolonged periods of favourable conditions for biomass accumulation. In contrast, intermediate latitudes such as Marseille (Calanques) and Bilbao exhibit more pronounced seasonality, with growth rates declining below ∼40% of maximum during winter and peaking at ∼60– 80% under summer conditions, while still maintaining relatively high annual performance (i.e., AGRs exceeding 9% day⁻¹ at both locations).

## 4. Discussion

Using the invasive macroalga *Rugulopteryx* as a case study, we show that combining physiological information with species distribution modelling can help interpret predictions of establishment potential in species that are not yet in equilibrium with their environment. Specifically, by comparing SDM-derived environmental thresholds with physiological limits as well as with physiological information from a closely related native species, we identified the ecological mechanism that is most likely to delimit the establishment range of *Rugulopteryx* in Europe. Beyond identifying areas where establishment may be possible, we demonstrate how physiological performance data can help predict where invasive species will most likely achieve high performance and may exert greater ecological impacts. Below, we discuss our findings and the corresponding implications for invasive species management.

### 4.1 Potential establishment range

We observed a considerable difference in the predicted northern establishment limit of *Rugulopteryx* depending on whether the SDMs were fitted using minimum temperature or temperature-driven annual growth as a predictor. In our study system, these variables represent winter survival and growth, respectively, two fundamental processes known to constrain ectotherm distributions in general, and seaweed distributions in particular (Breeman, 1988; Lüning, 1990; Andersen et al., 2015). Minimum temperature and annual growth strongly covary in the native range of *Rugulopteryx*, which prevented us from including both predictors in the same model. Such collinearity between predictors represents an important challenge for fitting SDMs, because it complicates the identification of which predictor, and hence, which underlying mechanism, is associated with specific range boundaries (Dormann et al., 2013). In the case of *Rugulopteryx*, this makes it difficult to infer from occurrence data alone whether establishment will be primarily constrained by lethal winter conditions or by temperatures supporting insufficient growth. Although collinearity between predictors does not immediately yield divergent geographic range predictions, complications arise when the relationship between such predictors changes in novel environments for which projections are made (Braunisch et al., 2013; Dormann et al., 2013). This issue is specifically relevant for invasive species, which are typically not yet in equilibrium with their invaded environment (Gallien et al., 2012). In this context, predictor choice has particularly large consequences, because the decoupling between collinear predictors in novel environments can lead to large discrepancies between model projections for these areas. For *Rugulopteryx* specifically, a change in the relationship between minimum temperature and annual growth in the invaded range resulted in the hybrid SDM producing a substantially more restrictive projection than the correlative model, reducing the projected northern establishment limit by more than 2,000 km (Fig. 5). As both models represent biologically relevant mechanisms that cannot be distinguished using occurrence data alone, alternative types of information, such as physiological data, are needed to assess the plausibility of these contrasting projections.

Assessing the thermal response of survival and growth for both *Rugulopteryx* and the native *Dictyota* allowed us to evaluate the plausibility of the predicted northern establishment range limits. The minimum temperature threshold identified by the correlative SDM (Fig. 5A, C) was consistent with the cold tolerance threshold determined in a lab experiment (4°C), providing physiological support for this prediction. In contrast, the hybrid model predicted a northern establishment limit corresponding to an average annual growth threshold of 7.1% day^-1^. As this threshold cannot be evaluated using the same growth data from which it was derived, a comparison with available growth data and distributional information from *Dictyota* was used as an ecological reference. *Dictyota* is able to sustain viable populations as far north as mid-Norway (Tronholm et al., 2010; Delva et al., 2026), where the species’ predicted annual growth based on the thermal performance curve obtained in this study is around 3% day⁻¹. This comparison suggests that the growth threshold identified by the hybrid model may be too restrictive and that the predicted average annual growth of *Rugulopteryx* at more northern locations may be sufficient to support population persistence, provided that other demographic processes (e.g., recruitment and overwinter survival) remain favourable. Taken together, these findings provide stronger support for the predictions made by the correlative model, suggesting that the northern establishment limit of *Rugulopteryx* is more likely to be constrained by winter survival than by insufficient thermal conditions for growth.

### 4.2 Projected growth performance and ecological impact

Predicting invasion risk based solely on establishment potential may not capture variation in the abundance or biomass of invasive species, which has often been linked to their ecological impact (Parker et al., 1999; Pearson et al., 2016; Bradley et al., 2019). In our study, we used temperature-dependent growth as an indicator for potential biomass accumulation and projected it onto monthly sea surface temperature data from across Europe to assess invasion risk. Our results revealed pronounced spatial variation in the projected seasonal growth of *Rugulopteryx*, with high year-round growth rates being projected throughout most of the Mediterranean basin and along the Atlantic coast up to northern Spain, while projected growth declined progressively north of the Bay of Biscay as monthly temperatures became colder. This resulted in highly seasonal dynamics in more northern locations, represented by increasingly shorter periods of active growth and lower peak growth rates during these periods. These results are consistent with other studies, which indicate that high performance, and hence, potentially high biomass accumulation, may only be achieved in parts of the establishment range (O’Neill et al., 2021). Importantly, in our study system, high performance was also predicted for parts of Europe that were predicted as unsuitable for the establishment of *Rugulopteryx*. Examples include parts of the eastern Mediterranean, where predicted absence is likely linked to high temperatures exceeding the species’ thermal tolerance during the hottest month, or in deeper regions, where establishment will be restricted by light availability. Because these environmental constraints are not captured by the performance data, our results highlight the importance of combining estimates of both establishment potential and performance when predicting invasion risk.

For many invaders, ecological impact is higher at temperatures that more closely match their thermal growth optima (Iacarella et al., 2015). This also implies that impacts decline as habitat conditions move further from this optimum (Ricciardi et al., 2013). Although this relationship has not been explicitly tested for invasive macrophytes, field studies have reported marked responses in invasion dynamics along thermal gradients for aquatic macrophyte and plant invaders (Wu et al., 2017; Muthukrishnan & Kalinowski, 2025). For *Rugulopteryx*, such relationship would suggest that the most severe impacts are likely to remain concentrated in southern Europe, where temperatures match the thermal growth optima of the species throughout most of the year, allowing for prolonged active growth and high growth rates. Indeed, field observations of extensive invasions (e.g., Ceuta or Marseille; (García-Gómez et al., 2020; Ruitton et al., 2021 but see also Marletta et al., 2024; Díaz-Tapia et al., 2025), coincide with projections of high growth rates throughout most of the year. The long-term introduction of *Rugulopteryx* in the Thau Lagoon provides a useful opportunity to test this hypothesis. There, projections of seasonal growth indicate similar response to that projected at the northern limit of the native distribution (Fig. S3, Fig. S4) with peak growth occurring in spring and summer and reduced growth during winter (November to March). In this lagoon, sea surface temperatures frequently decrease to around 5°C during the coldest month (Pigeon et al., 2026). Despite being present for more than 20 years, *Rugulopteryx* has not exhibited invasive behaviour. Instead, established populations persist as low-density strands that undergo strong winter regressions (Ruitton et al., 2021)

Differences in thermal optima can further create a spatial shift in competitive advantage. In the Strait of Gibraltar, the dominance of *Rugulopteryx* over native macroalgae has been linked to its rapid growth, itself enabled by thermal conditions that remain close to the species’ growth optimum for most of the year (Hwang et al., 2009; García-Gómez et al., 2020; Mercado et al., 2022). Our results suggest that this advantage may progressively weaken toward higher latitudes as environmental conditions become less favourable for high growth and biomass accumulation. Such shifts in competitive ability have been previously documented for invasive species, with ecological dominance varying along environmental gradients as conditions become more or less favourable for invader performance (Thiele et al., 2010; Seipel et al., 2016; Pulzatto et al., 2019). In this context, native species that are more cold-tolerant, such as *Dictyota,* may represent an additional constraint on *Rugulopteryx* as it spreads towards the northern edge of its potential establishment range.

Growth serves as a useful proxy for local biomass accumulation and potential ecological impact in our study system, but traits governing abundance and impact are likely to differ among invaders. In marine fishes for instance, ecological impacts are often better explained by behavioural traits such as feeding efficiency or plasticity (Green et al., 2011; Vergés et al., 2014) whereas for terrestrial plants, these may be more strongly driven by chemical defense or drought tolerance (Keane & Crawley, 2002; Callaway & Ridenour, 2004). For *Rugulopteryx*, the mechanisms driving ecological impacts remain only partially understood. Rapid growth is frequently cited as a key component of the species’ invasive success, but high propagule pressure generated through extensive vegetative proliferation is also likely to play an important role in local recruitment and abundance (Altamirano et al., 2019; Rosas-Guerrero et al., 2026). Likewise, the formation and persistence of dense stands may also be facilitated by the production of effective chemical defences that may reduce herbivory in the invaded range (Casal-Porras et al., 2021). Consequently, careful consideration should be given to trait selection when using physiological performance to approximate invasion impacts. Ultimately, it will likely be the interaction between multiple fitness-related traits and the local environment that determines realized local abundance and ecological impacts.

### 4.3 Management recommendations

Importantly, our projections suggest that establishment potential and high performance can be spatially decoupled, implying that management priorities should differ geographically. In northern Europe, where the species has not yet established but suitable habitat appears widespread, prevention should focus on pathway management, surveillance and early detection around high-risk introduction points such as major commercial ports and other hubs of maritime activity. In contrast, parts of the central and eastern Mediterranean face both high establishment potential and prolonged growth. These regions should continue surveillance alongside measures to reduce secondary spread and mitigate ecological and socio-economic impacts where feasible. Given the limited evidence for effective control of established *Rugulopteryx* populations, our results highlight the importance of acting before widespread establishment. Building upon the experience gained in heavily invaded regions (MITECO 2022), coordinated management strategies including biomass removal or valorisation could help mitigate ecological and socio-economic impacts. The recent inclusion of *Rugulopteryx okamurae* on the European Union list of Invasive Alien Species of Union concern (European Union (EU), 2022) provides a timely legal framework for implementing pathway management and surveillance.

Using the macroalgae invader *Rugulopteryx okamurae* as a case study, we show that areas vulnerable to establishment in Europe are not necessarily those where performance, and hence, potential ecological impacts will be greatest. Explicitly separating these two dimensions can help align prevention, surveillance and long-term mitigation with the stage and expected severity of invasion. Our framework identifies where *Rugulopteryx* is most likely to establish and where favourable thermal conditions may support rapid growth. Conservation prioritisation additionally requires information on the spatial distribution of vulnerable habitats and species. Future work should focus on overlaying the results of the physiologically informed SDMs with maps of conservation assets (e.g. kelp forests, *Posidonia oceanica* meadows, coralligenous and rocky reef communities or marine protected areas in general). This would enable managers to identify locations where invasion pressure and conservation value coincide, thereby maximising the effectiveness of surveillance and intervention and delivering the greatest conservation benefit.

## Author Contributions

**Lauren Vapillon**: methodology, formal analysis, visualization, writing – original draft, writing - review and editing. **Soria Delva**: conceptualization, methodology, writing – original draft, writing – review and editing. **María Bonafont Castelles**: methodology, data curation, writing – original draft. **Jorge Assis**: methodology, writing – review and editing. **Diederik Strubbe**: writing – review and editing. **Tim Adriaens**: writing – review and editing.: **Olivier De Clerck**: conceptualization, data curation, funding acquisition, supervision, writing – review and editing. **Sofie Vranken**: conceptualization, supervision, writing – original draft, writing – review and editing.

