## Supplementary material for "Beyond establishment: incorporating physiological performance into predictions of invasion risk": Fig. S

Supplementary Materials

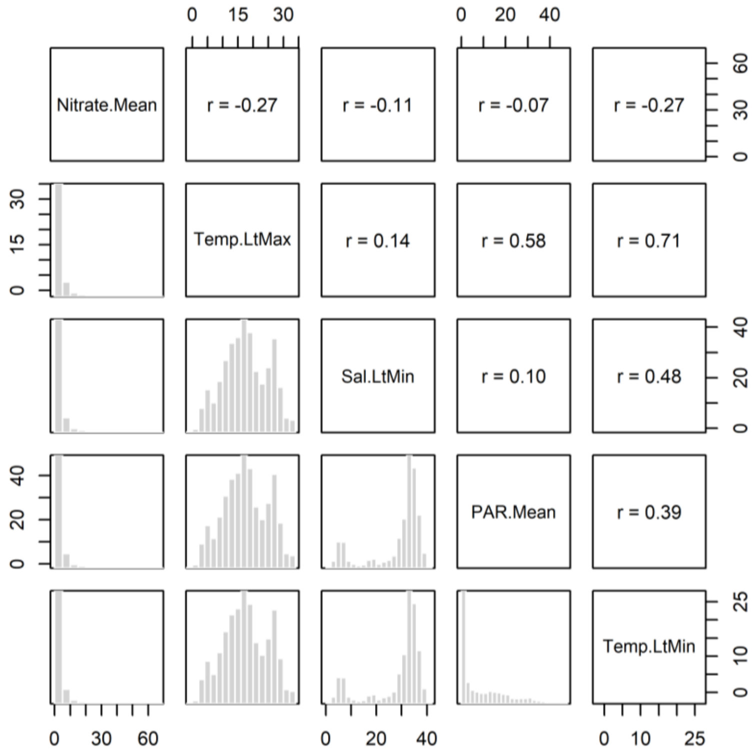


*Fig. S1. Pearson's correlation (r) calculated between each pair of variables. In the Hybrid approach, Annual.Gowth is replacing Temp.Lt.Min and is correlated to Temp.Lt.Max with r=0.80.*

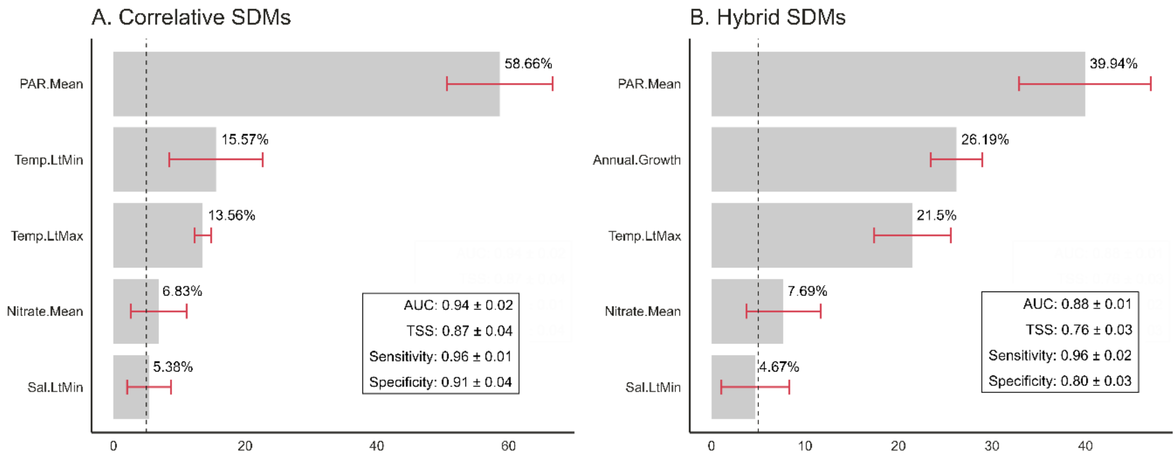


*Fig. S2. Mean relative contribution (%) of environmental predictors used to model likelihood suitability across all 5 replicates of the A) Correlative and B) Hybrid models. Red error bars represent the standard deviation across runs, and dashed lines depict contributions >5%. Model performances (AUC, TSS) are reported for each model type as the mean + sd across replicates.*

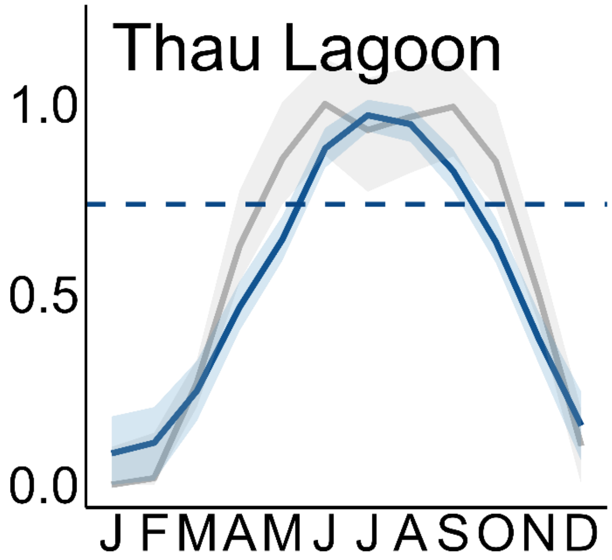


*Fig. S3. Projected seasonal growth of Rugulopteryx (coloured line) and Dictyota (grey line) at Thau Lagoon (France), standardized to the maximum predicted growth rate of each species (range: 0–1) as described by the Lobry model (Lobry, 1991). Shaded ribbons represent the 95% confidence intervals obtained by bootstrap resampling (n = 1000). Horizontal dashed lines depict the annual growth rates of Rugulopteryx at the site.*

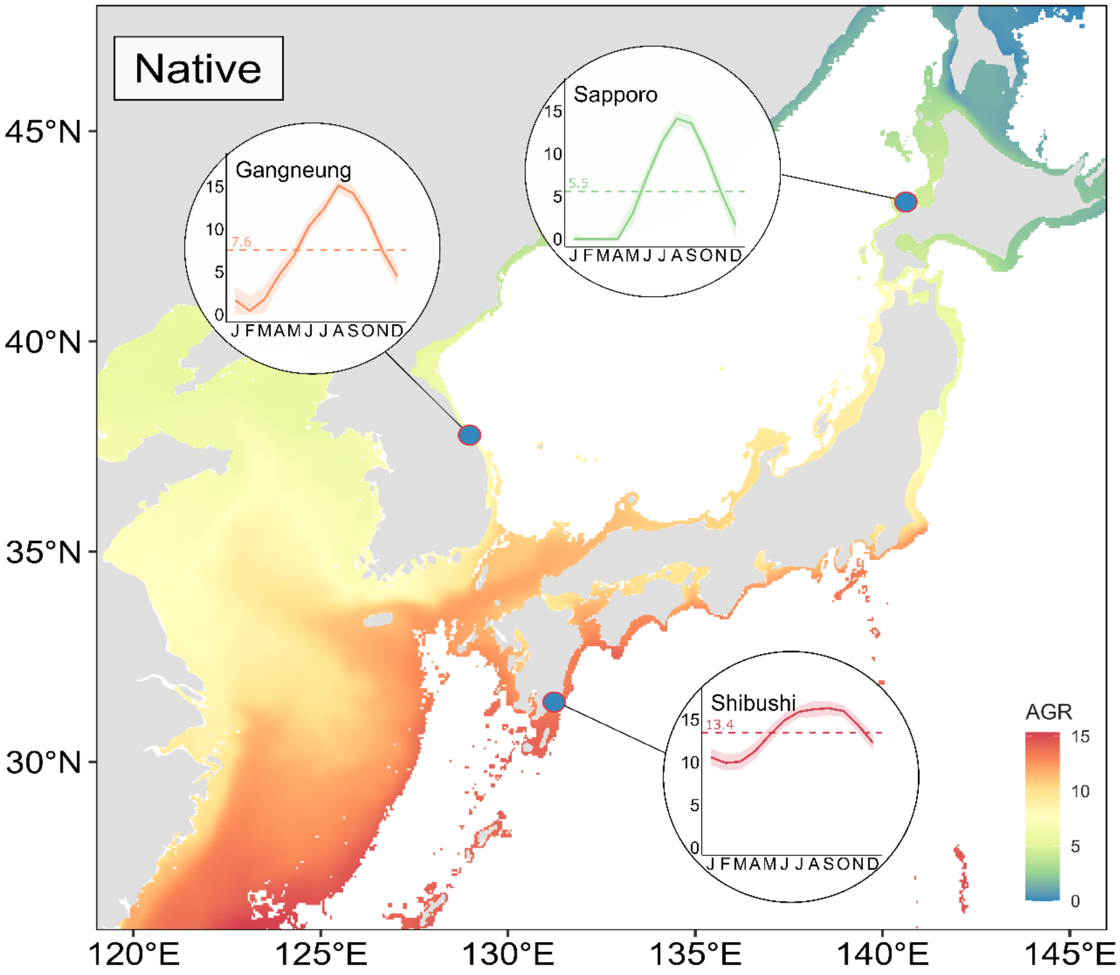


*Fig. S4. Annual growth rates (AGR) projected in the native range of the species. Bubbles depict the projected seasonal growth of Rugulopteryx (coloured lines), expressed as absolute specific growth rates (% day $^{-1}$), as described by the Lobry model (Lobry, 1991). Shaded ribbons represent the 95% confidence intervals obtained by bootstrap resampling (n = 1000). Horizontal dashed lines depict the annual growth rates of Rugulopteryx at the respective sites.*

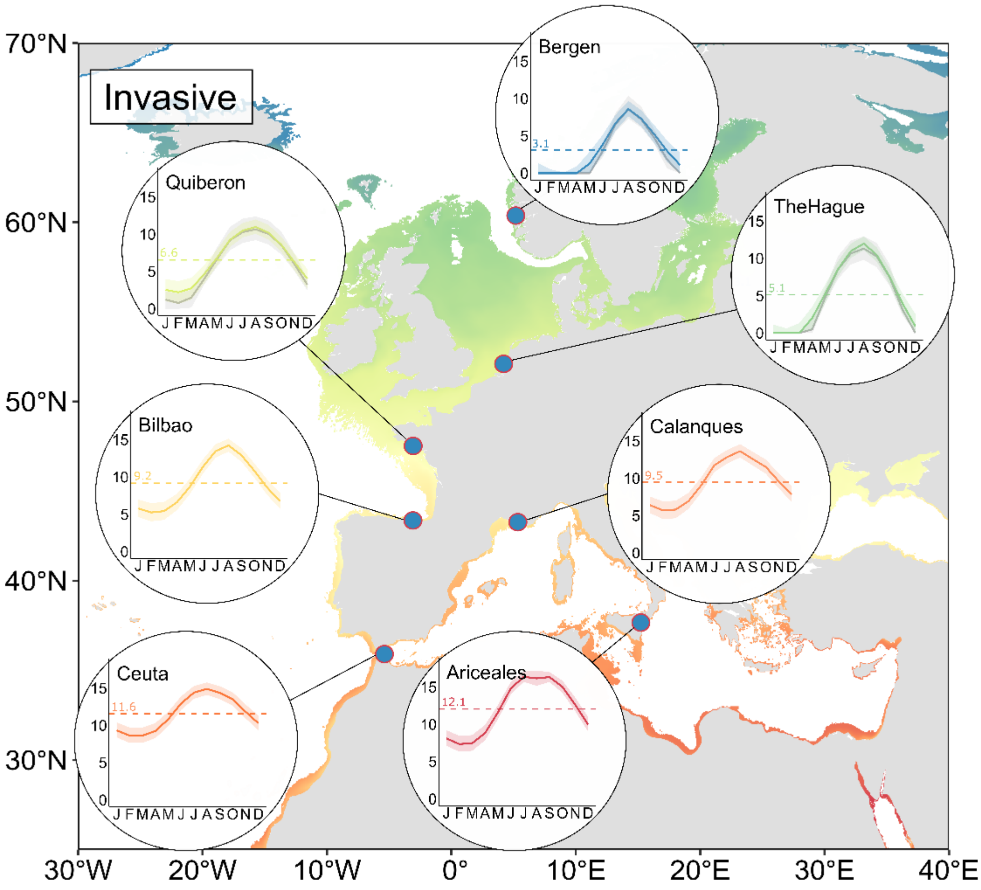


Fig. S5. *Annual growth rates (AGR) projected in the invasive range of the species. Bubbles depict the projected seasonal growth of Rugulopteryx (coloured lines) and Dictyota (grey lines; projected at Quiberon, The Hague and Bergen), expressed as absolute specific growth rates (% day* $^{-1}$*), as described by the Lobry model (Lobry, 1991). Shaded ribbons represent the 95% confidence intervals obtained by bootstrap resampling (n = 1000). Horizontal dashed lines depict the annual growth rates of Rugulopteryx at the respective sites.*

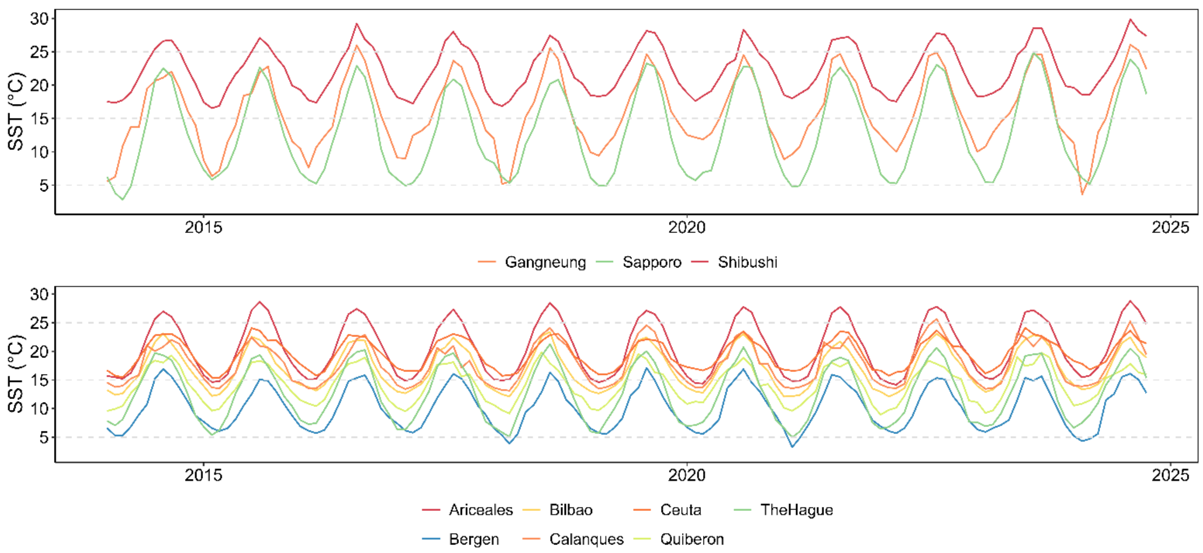


*Fig. S6. Sea Surface Temperature (SST) Time series at all 10 locations investigated across the A) Native and B) Invasive range of the species.*

*Table S1. Minimum, Maximum, and Long-term averages of Sea Surface Temperature (SST) at all 10 locations investigated across the Native and Invasive range of the species.*

| | Sites | Min | Max | LtMin | LtMax |
|---|---|---|---|---|---|
| Invasive | Calanques | 12.6 | 28.7 | 13.2 | 26.0 |
| | Ceuta | 14.7 | 25.4 | 15.7 | 24.2 |
| | Ariceales | 13.8 | 29.3 | 14.5 | 28.4 |
| | Bilbao | 11.4 | 24.8 | 12.1 | 23.8 |
| | Quiberon | 6.8 | 21.2 | 8.4 | 20.4 |
| | TheHague | 3.7 | 22.1 | 5.2 | 21.0 |
| | Bergen | 2.4 | 19.8 | 4.7 | 17.3 |
| Native | Sapporo | 2.0 | 26.5 | 4.5 | 24.1 |
| | Shibushi | 15.7 | 31.0 | 17.0 | 29.4 |
| | Gangneung | 2.5 | 28.1 | 6.1 | 26.0 |
